# EEG signal diversity during propofol sedation: an increase in sedated but responsive, a decrease in sedated and unresponsive subjects

**DOI:** 10.1101/444281

**Authors:** Michał Bola, Paweł Orłowski, Martyna Płomecka, Artur Marchewka

**Affiliations:** Laboratory of Brain Imaging, Nencki Institute of Experimental Biology of Polish Academy of Sciences, Warsaw, Poland; Institute of Philosophy, University of Warsaw, Warsaw, Poland; Faculty of Mathematics, Informatics and Mechanics, University of Warsaw, Warsaw, Poland

**Keywords:** consciousness, propofol, EEG, complexity, diversity

## Abstract

Transitions between wakefulness and anesthesia are accompanied by profound changes in brain functioning. A key challenge is thus to disentangle neuronal mechanisms specific to loss and recovery of consciousness, from more general effects that are not directly related to the capacity for conscious experience. Measures of neuronal diversity have been recently proposed to constitute a robust correlate of the global states of consciousness. In the present study we investigated whether EEG signal diversity is indeed related to behavioral responsiveness during propofol sedation, or rather to the general drug-related effects. To this end, we reanalyzed data collected from 20 subjects sedated with propofol. Based on the responsiveness to auditory stimuli all subjects were subdivided into two subgroups - *responsive* (n = 13), who remains awake throughout the experiment, and *drowsy* (n = 7), who becomes unresponsive during moderate sedation. Resting state EEG recorded during wakefulness and sedation was characterized by the Mean Information Gain (MIG) and Fluctuation complexity (FC) - information-theory measures estimating signal diversity or complexity, respectively. The main finding is that the *drowsy* group exhibited a decrease in diversity during sedation but, unexpectedly, the *responsive* group exhibited a robust increase in diversity (ANOVA *group* x *state* interaction: F(3) = 7.81, p < 0.001; BF_10_ > 197). However, signal complexity neither differentiated the subgroups, nor decreased reliably during sedation (t-test wake vs. moderate sedation: t(19) = 2.57, p = 0.092; BF_10_ = 3.08). Further, we show that a change in signal diversity is negatively correlated with a delta power change (r = −0.62, p = 0.002), and positively correlated with a beta power change (r = 0.84, p < 0.001). Finally, we show that MIG behaves in a qualitatively similar manner to Lempel-Ziv – another diversity measures used in several recent studies. Overall, we revealed that propofol sedation is initially related to an increase in EEG signal diversity, and that only upon loss of responsiveness EEG diversity decreases. The qualitatively different pattern of changes in the *responsive* and *drowsy* groups makes EEG diversity a robust indirect index of responsiveness and, presumably, consciousness.

## Introduction

What are the neuronal mechanisms of consciousness? One of the main hypotheses is that the capacity for conscious experience relies on the brain’s ability to generate a large repertoire of functional states (Seth et al., 2006; Tononi and Edelman, 1998). Importantly, the temporal diversity of brain states needs to be balanced between order and disorder, as both insufficient temporal diversity (i.e. dwelling in the same state for long time periods) and excessive diversity (i.e. chaotic transitions between states) will likely cause alterations in the global state of consciousness.

Evidence supporting this hypothesis has been provided by studies using a perturbational approach, which involves activating a local brain region by applying a TMS pulse and recording the spatio-temporal spread of activity using EEG. It has been shown that the EEG response is complex during conscious states, as it is spatially integrated and at the same time temporally differentiated, but becomes simple and stereotypical during NREM sleep (Massimini et al., 2005). To quantify the complexity of brain response the Perturbational Complexity Index (PCI) has been developed, which is based on the estimation of Lempel-Ziv complexity of the TMS-evoked EEG response (Casali et al., 2013). PCI indeed indicates high complexity of brain response during conscious wakefulness, but a robust decrease is observed during NREM sleep (Pigorini et al., 2015) or general anesthesia (Ferrarelli et al., 2010; Sarasso et al., 2015). Due to an exceptional sensitivity PCI has been subsequently applied in the clinical context to successfully classify patients with disorders of consciousness (Casarotto et al., 2016; Rosanova et al., 2012). Promising results of the PCI studies encouraged researchers to investigate similar properties in the spontaneous brain activity recorded during different states of consciousness. Similarly to PCI, diversity of spontaneous EEG signals is high during wakefulness and drops during propofol anesthesia or NREM sleep (Schartner et al., 2015, 2017; Andrillon et al., 2016).

Based on these findings brain signal diversity has been interpreted as a correlate of consciousness. However, conscious and unconscious states differ not only with respect to the presence or absence of conscious experiences, but also multiple other physiological and mental processes. Thus, a key challenge is to disentangle these effects in order to distil the neuronal mechanisms specific to consciousness. Several different approaches have been employed in order to address this problem in the context of diversity measures. First, one can assume that the genuine mechanisms of loss and recovery of consciousness are independent of the specific cause for a transition between the states, and thus a given measure should be tested in a range of unconscious states. Indeed, both PCI and spontaneous diversity are associated with loss and recovery of consciousness during sleep, anesthesia caused by different pharmacological agents, and in brain injury patients (e.g. Casali et al., 2013). Second, one might aim to dissociate behavioral responsiveness from consciousness by studying conscious but unresponsive states. Again, both PCI and spontaneous diversity remain high during REM sleep and ketamine sedation, which indicates they are sensitive to experiences in the form of dreams and hallucinations, irrespective of behavioral responsiveness (Sarasso et al., 2015; Schartner et al., 2017). Third, if a given measure indeed reflects quality of conscious experience one can aim to manipulate some aspects of phenomenology to see if the tested measure is sensitive to such manipulations. Interestingly, recent studies reported increases in spontaneous diversity of brain states during psychedelic states caused by LSD or ketamine (Schartner et al., 2017b; Wang et al., 2017; Farnes et al., 2019; Tagliazucchi et al., 2014), or by stroboscopic light stimulation (Schwartzman et al., 2019). This suggests that not only loss of consciousness, but also alterations of the phenomenology towards a more psychedelic type are reflected by changes in brain signal diversity.

Yet another possibility to test whether a given measure is indeed a candidate for a correlate of consciousness is by taking advantage of the between-subjects variability in susceptibility to anesthetics (Palanca et al., 2009) and comparing brain activity observed in subjects who are sedated and unconscious, with subjects who are sedated with similar anesthetic doses but remain conscious. If a given measure is indeed sensitive to consciousness, then it would change in the unconscious group only, but if a measure is rather related to the general effects of an anesthetic, then a change would be observed in both groups.

In the present study we adopted such an approach to investigate whether signal diversity is indeed related to behavioral responsiveness and, presumably, consciousness during propofol sedation. We reanalyzed data from an experiment conducted by Chennu and colleagues (2016), in which all subjects were sedated with similar doses of propofol, but based on their responsiveness to auditory stimuli they could be subdivided into two subgroups: sedated but responsive, and sedated and unresponsive. Resting-state EEG recorded during wakefulness and sedation was characterized in terms of two information-theory measures: the Mean Information Gain (MIG), which measures *diversity* understood as randomness or entropy of the data, and Fluctuation Complexity (FC), which is a measure of *complexity* understood as the state intermediate between order and disorder (**Fig. 1**). We hypothesized to observe a decrease of diversity and complexity during sedation, but only in the unresponsive group.

**Figure 1.**
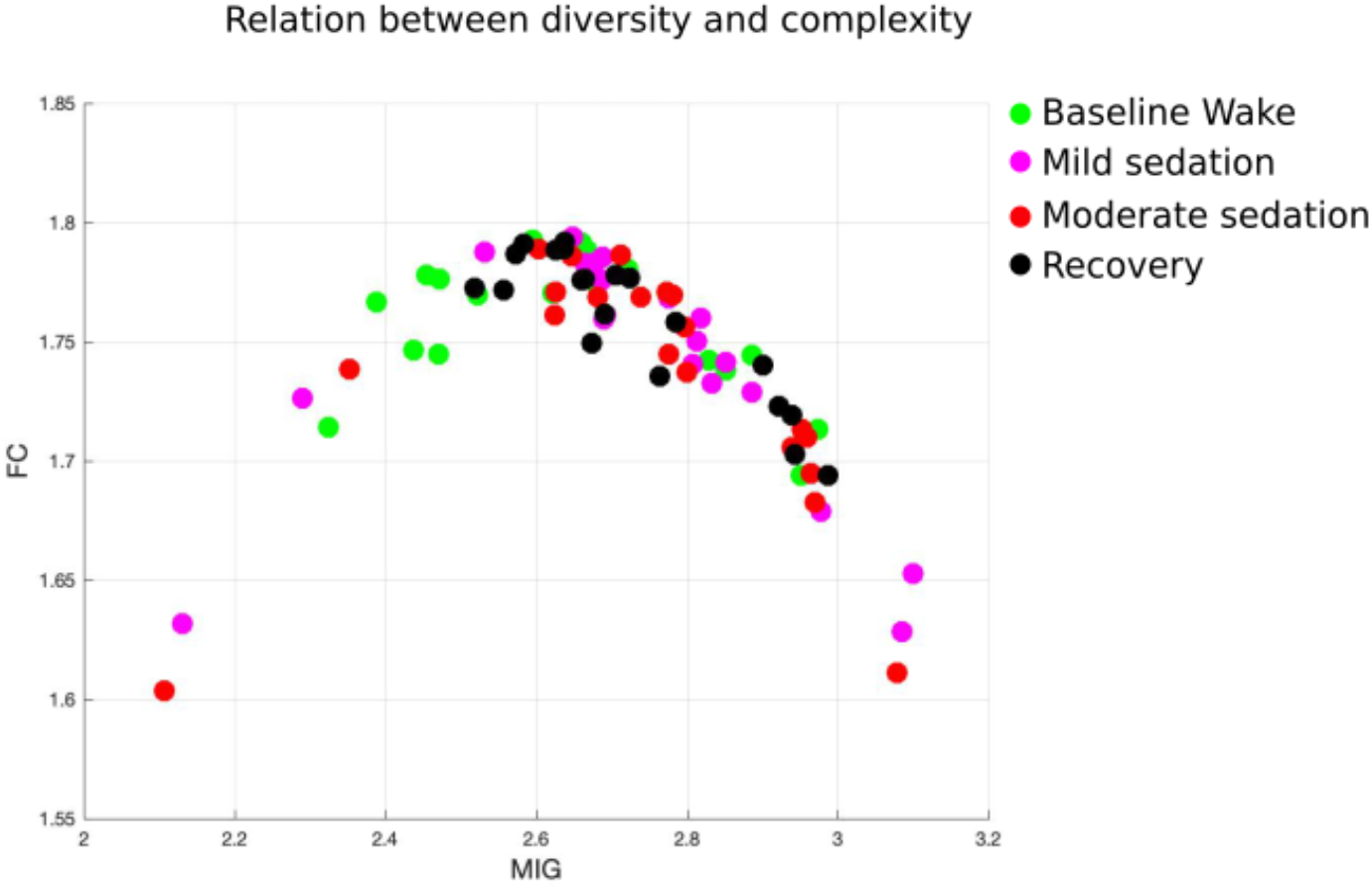
A relation between complexity (FC) and diversity (MIG). Each dot represents a value calculated from each state and subject (altogether 20 x 4 = 80 data-points). The data show an inverted “U” shape, as complexity peaks for signals of intermediate diversity (between order and disorder).

## Methods

### Subjects and data collection

In the present study we reanalyzed data first published by Chennu et al. (2016). The procedures regarding data collection and preprocessing are briefly described below and the original paper should be consulted for additional details. Ethical approval for the study was provided by the local ethics committee and all subjects provided a written informed consent. The data are accessible from https://www.repository.cam.ac.uk/handle/1810/252736.

Data collected from 20 healthy subjects (11 females; mean age = 30.85, SD = 10.98) were used in the present analysis. Data were collected in four states: wakeful baseline, mild sedation, moderate sedation, and recovery. During each state subjects first performed an auditory discrimination task (for 3 to 5 minutes), and afterwards 7 minutes of eyes-closed resting-state EEG was recorded. States of mild and moderate sedation were obtained by performing a target-controlled infusion of propofol with a computerized syringe driver. The targeted blood plasma levels of propofol were 0.6 mcg/ml for mild sedation and 1.2 mcg/ml for moderate sedation. At each targeted level a period of 10 minutes was allowed to obtain a steady propofol concentration and only afterwards behavioral and EEG measurements were performed. Blood samples were taken in each state to evaluate the actual propofol concentration in blood.

To evaluate subjects’ responsiveness in each state they were asked to perform a simple auditory discrimination task. Specifically, to indicate, by pressing on of two buttons, whether a presented binaurally stimulus was a broadband noise (“noise”) or a harmonic complex with a 150 Hz fundamental frequency (“buzz”). 40 stimuli (20 of each kind) were presented in a random order with a mean interstimulus interval of 3 s. A *hit-rate* was defined as a proportion of correct responses. Binominal modelling of hit-rates obtained during baseline wakefulness and moderate sedation was used to classify subject into either drowsy or responsive subgroups (details of the procedure can be found in: Chennu et al., 2016).

In each state approximately 7 minutes of eyes-closed resting-state EEG was recorded with high-density 128 electrodes caps and the Net Amps 300 amplifier (Electrical Geodesic Inc., Eugene, Oregon, USA). Offline preprocessing was conducted with EEGlab (Delorme and Makeig, 2004). First, 37 channels placed over neck, cheeks, and forehead were excluded and only data from 91 channels covering the scalp area were retained and further analyzed. EEG signals were filtered between 0.5 Hz and 45 Hz and segmented into 10-seconds long epochs. Mean value was subtracted from signal from each channel in an epoch. Semi-automatic procedure was used to remove noisy epochs and channels. The mean ± SD number of retained epochs was: baseline wakefulness - 38 ± 5; mild sedation - 39 ± 4; moderate sedation - 38 ± 4; recovery - 40 ± 2 and did not differ significantly across states. Missing channels were interpolated using the spherical spline interpolation algorithm. As the last step, the data were re-referenced to the average of all channels.

The EEG data were published in the already preprocessed form and we did not apply any further data transformations before calculating the measures described below.

### Diversity and complexity analysis

#### a) Mean information Gain (*MIG*)

The Mean Information Gain (*MIG*) is one of the information theory-based measures. *MIG* is a measure of diversity, as it exhibits maximum values for random signals. The details and a formal definition of *MIG* can be found in: Bates and Shepard (1993) and Wackerbauer et al. (1994). An application of *MIG* to EEG data was described by Wang et al. (2017) and by Bola et al. (under review), and here we followed the same procedure.

Briefly, to calculate *MIG* a signal has to be binarized. Here, for each epoch and channel separately, a median value was used as a threshold, and “1” was assigned to the time-points exceeding the median value, while “0” was assigned to the time-points that were below or at the median value. Then binary series were partitioned into “words” of *L* length, where *L* is the “window length’ or ‘information length’, and each “word” reflects the state of a system at a given time-point. Using *L* = 4, as in the main analysis reported here, the possible states/words were 0000, 0001, 0010, 0100, 1000, 0011,0110, 1100, 1010, 0101, 1001, 0111, 1011, 1101, 1110, 1111. A transition to a next state occurs by a shifting a window forward by one symbol.

Calculating *MIG* starts with the Shannon information content. We define the probability *p_i_* that the system is currently in a state ‘*i*’ by dividing the number of occurrences of a state ‘*i*’ by the total number of occurrences^19^. Based on the probabilities of observing transition from state *i* to *j*, defined as *p_ij_*, we calculate the transition probability 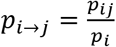. In the same way, the reverse conditional probability can be obtained as 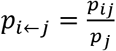.

The net information gain can be written as 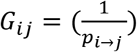, and by averaging the gains over all transitions the *MIG* is estimated:

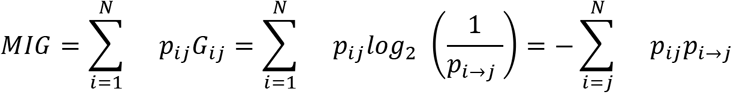

#### b) Fluctuation Complexity (*FC*)

Another example of an information theory-based measure is Fluctuation Complexity (*FC*). The details and a formal definition of *FC* can be found in: Bates and Shepard (1993) and Wackerbauer et al. (1994). While applying *FC* to EEG data we again followed the procedure described in Wang et al. (2017) and Bola et al. (under review). Unlike *MIG, FC* does not consider a random signal to be complex. In the computation of *FC*, the EEG signals were binarized in the same way as described for *MIG*.

While shifting from one state to another, the newly observed symbol contributes to a gain of information, defined as 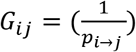. At the same time we forget the first symbol from the previous word. The loss of information can be expressed as 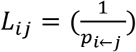.

In average for the whole symbol sequence, information gain and loss eliminate each other. The fluctuation complexity is the mean square deviation of the net information gain (i.e. the differences between information gain and loss):

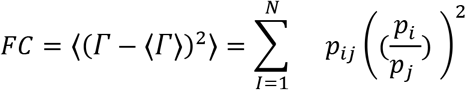

The more this balance of information gain and loss is fluctuating in the given EEG time series the more complex is the signal in the sense of the fluctuation complexity.

#### c) Lempel-Ziv (*LZs* and *LZc*)

*MIG* and *FC* have used in recent studies to evaluate diversity and complexity of EEG signals (Wang et al., 2017). But majority of previous studies have employed the Lempel-Ziv algorithm to estimate EEG signal diversity (Casali et al., 2013; Schartner et al., 2015). Therefore, for consistency with the previous studies in the present analysis we used two version of the Lemepl-Ziv measures, single-channel Lempel-Ziv (*LZs*), which captures temporal variability, and multi-channels Lempel-Ziv (LZc), which captures spatio-temporal variability. Similarly to *MIG*, Lempel-Ziv is a measure of diversity, as it reaches maximum values for random signals. We used an implementation of the method described by Schartner et al. (2017). Results of the *LZs* and *LZc* analysis are presented in the supplementary materials.

To estimate *LZs* EEG signal from each channel and epoch was assessed independently by demeaning (within 10 s long segment), dividing by standard deviation, and removing a linear trend. Hilbert transform was used to estimate the envelope of a signal (an absolute value of the analytic signal) and the mean of the envelope was used as a threshold to binarize the signal, i.e. “1” was assigned to time-points exceeding the mean value, while “0” was assigned to measurements that were below or at the mean value. The binarized signal was then segmented into blocks using the encoding step of the Lempel-Ziv compression algorithm. The number of blocks is the raw *LZs* score for the segment. Next the raw *LZs* value was normalized by the raw *LZs* value obtained from the same binary signal shuffled in time. Thus, the final (normalized) *LZs* score varies between 0 and 1 indicating, respectively, minimally and maximally diverse signals.

To estimate *LZc* the 64 time series (one for each EEG channel) from a 10 s segment of data are binarized as described above for *LZs*, then concatenated observation-by-observation into one binary string such that the first 64 digits of that string are the observations of the 64 channels at time step 1, the next 64 are those at time step 2, etc. The diversity of this binary string is then assessed in the same way as described for *LZs* above.

#### d) Spectral power density

The spectral power density of the signals was estimated using the EEGlab *spectopo* function, which returns power density estimates transformed logarithmically and analyzed as 10log^10^(μV^2^/Hz). For each subject and condition, the power spectra were averaged over all electrodes and next averaged within the following classic frequency ranges: delta 1-4 Hz; theta 4-8 Hz; alpha 8-15 Hz; beta 15-25 Hz.

### Statistical analysis

All values are presented as Mean ± Standard Error of the Mean (SEM), unless stated otherwise. Statistical analyses were conducted in JASP software (Wagenmakers et al., 2018) and in Matlab. The performed tests involved two-way repeated-measures ANOVA, with within-subjects factor *state* (4 levels: wakeful baseline, mild sedation, moderate sedation, recovery) and between-subjects factor *group* (2 levels: drowsy, responsive). Subsequent post-hoc tests were computed, either paired-samples t-tests to compare between states, or independent-samples t-tests to compare between groups. For each measure post-hoc tests were corrected for multiple comparisons (6 when comparing between the states, 4 when comparing between the subgroups within each state) using the Bonferroni-Holm procedure (Holm, 1979).

The traditional null-hypothesis significance testing approach was complemented with Bayesian statistics in order to enable testing for the lack of differences between variables. The Bayes Factor (BF_10_) is defined as the ratio of the probability of observing the data given the alternative hypothesis is true, to the probability of observing the data given the null hypothesis is true. Thus BF_10_ provides a continuous measure of evidential support and is typically interpreted according to Lee and Wagenmakers (2013): BF_10_ < 0.1 indicates strong evidence in favor of the null hypothesis (i.e. lack of an effect); 0.1 < BF_10_ < 0.33 indicates moderate evidence in favor of the null hypothesis, 0.33 < BF_10_ < 3 indicates inconclusive data, 3 < BF_10_ < 10 indicates moderate evidence in favor of the alternative hypothesis, BF_10_ > 10 indicates strong evidence in favor of the alternative hypothesis,. Bayesian equivalents of a two-way ANOVA and t-test were calculated using a medium prior scale (Cauchy scale: .707).

In an additional analysis we aimed to test whether between-group differences in MIG observed during moderate sedation can be explained by differences in propofol concentration. We thus conducted a one-way ANOVA with a *group* factor, which in this case is equivalent to an independent-samples t-test, and included a propofol concentration as a covariate.

Finally, in order to test the ability of the measures used (diversity, complexity, spectral power) to distinguish between *responsive* and *drowsy* groups we conducted we conducted the Area Under the receiver operating characteristic Curve (AUC). Specifically, a logistic regression model with binominal distribution was fitted to the data (*fitglm* matlab function). Next the Receiver Operating Characteristic curve was calculated and the area under the curve was estimated (*perfcurve* function). Generally, AUC = 0.5 indicates that a given measure cannot discriminate two groups above chance level, whereas AUC=0 or 1 means that a threshold exists which divides all data-points correctly into two classes (i.e. allows perfect classification). Typically, measures characterized by AUC < 0.25 or > 0.75 are considered to have good discriminative power.

## Results

### Behavioral results and propofol concentration

In the present study we reanalyzed a data-set published by Chennu et al. (2016). In their study 20 subjects were sedated with propofol and data were acquired in four states: baseline wakefulness, mild sedation, moderate sedation, and recovery. In each state subjects performed an auditory discrimination task to assess their responsiveness and next resting-state EEG was recorded. The profiles of hit rates, defined as a proportion of correct responses, indicate that subjects can be subdivided into two subgroups - *responsive* (n = 13), who remains awake throughout the experiment, and *drowsy* (n = 7), who becomes unresponsive during moderate sedation (**Fig. 2**). Comparing both groups in terms of propofol concentration in blood we found a significant *state* x *group* interaction (F(2) = 4.68, p = 0.016; BF_10_ = 3.33). Specifically, there was no difference between groups during mild sedation (0.41 ± 0.04 vs. 0.50 ± 0.08 mcg/ml; t(18) = 1.05, p = 0.30; BF_10_ = 0.59) and recovery (0.27 ± 0.02 vs. 0.32 ± 0.03; t(18) = 1.37, p = 0.18; BF_10_ = 0.77), but during moderate sedation the *drowsy* group had a higher in-blood propofol concentration (0.79 ± 0.05 vs. 1.09 ± 0.12; t(18) = 2.60, p = 0.018; BF_10_ = 3.37).

**Figure 2.**
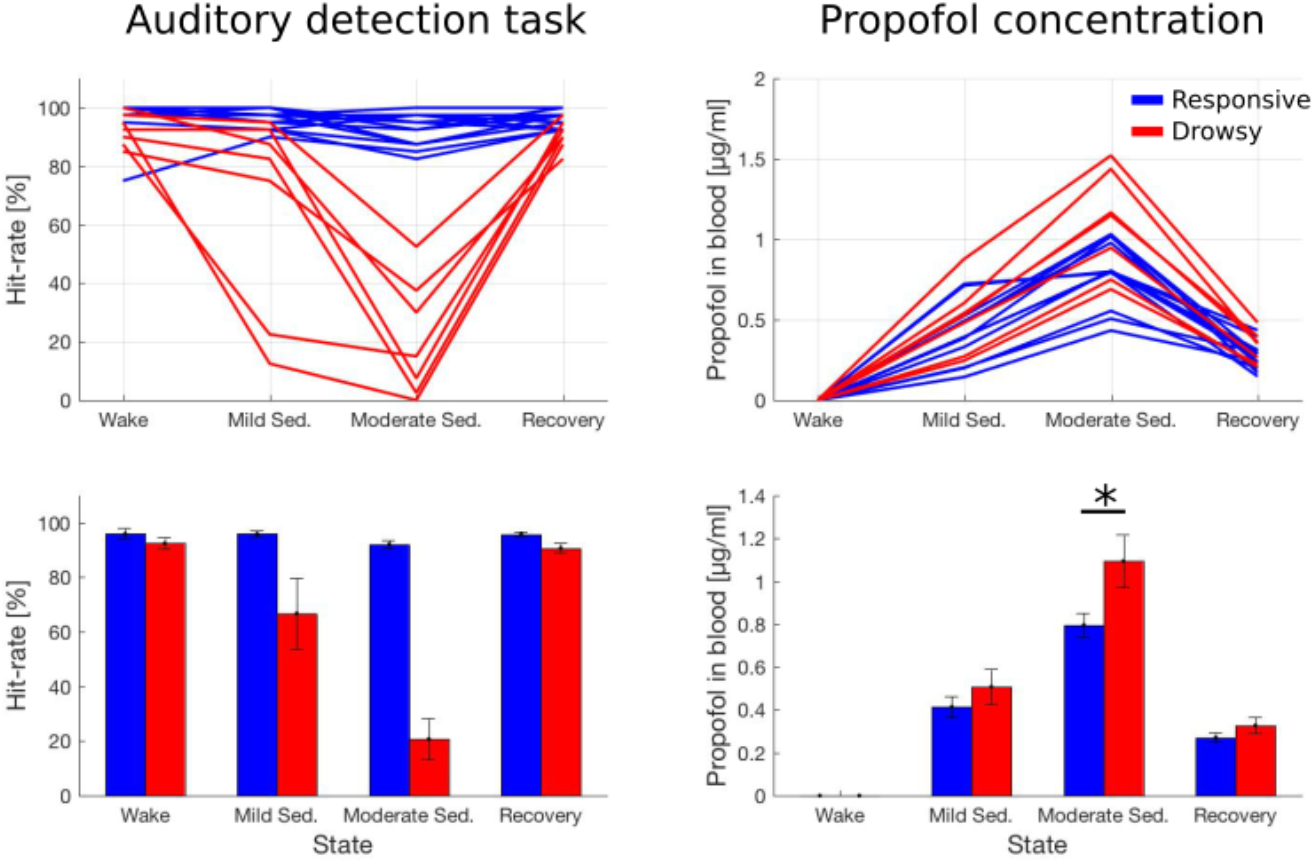
Hit-rates during an auditory detection task (left panels) and propofol concentration estimated from blood samples taken during the experiment (right panels). Upper panels present data of individual subjects from both groups, while bar-plots in lower panels present mean ± SEM of the same data. Statistical analysis of hit-rates was not conducted in this study, and in case of propofol concentration only comparisons between groups (but within states) were conducted. * - p < 0.05.

### Changes in diversity and complexity

The recorded resting-state EEG signals were characterized in terms of diversity and complexity using two information theory measures (Bates and Shepard, 1993; **Fig. 1**). Diversity, understood as randomness of the data, was estimated by the Mean Information Gain (MIG). Statistical analysis of MIG indicates no main effect of *group* (F(1) = 1.8, p = 0.19; BF_10_ = 0.80) and no effect of *state* (F(3) = 2.59, p = 0.062; BF_10_ = 1.27), but a significant *group x state* interaction was found (F(3) = 7.81, p < 0.001; BF_10_ = 197; **Fig. 3**). Subsequent post-hoc tests indicate that, with respect to the wakeful baseline, MIG increases during mild sedation (t(19) = 3.34, p = 0.020; BF_10_ = 12.7) and recovery (t(19) = 3.15, p = 0.026; BF_10_ = 8.83), but not during moderate sedation (t(19) = 2.47, p = 0.093; BF_10_ = 2.56). Importantly, lack of the significant effect during moderate sedation is due to a different pattern of changes in both subgroups, which is confirmed by a significant difference between groups during moderate sedation (t(18) = 3.24, p = 0.005; BF_10_ = 9.20) but not during other states (all p > 0.10; all 0.3 < BF_10_ < 1). Specifically, the *drowsy* group exhibits a decrease in MIG during moderate sedation (with respect to baseline) but the *responsive* group exhibits an increase. The topographic maps indicate that the increase of MIG in the responsive group is most prominent over the frontal and central midline regions, whereas the decrease in the drowsy group is greatest laterally over central regions (**Fig. 4**).

**Figure 3.**
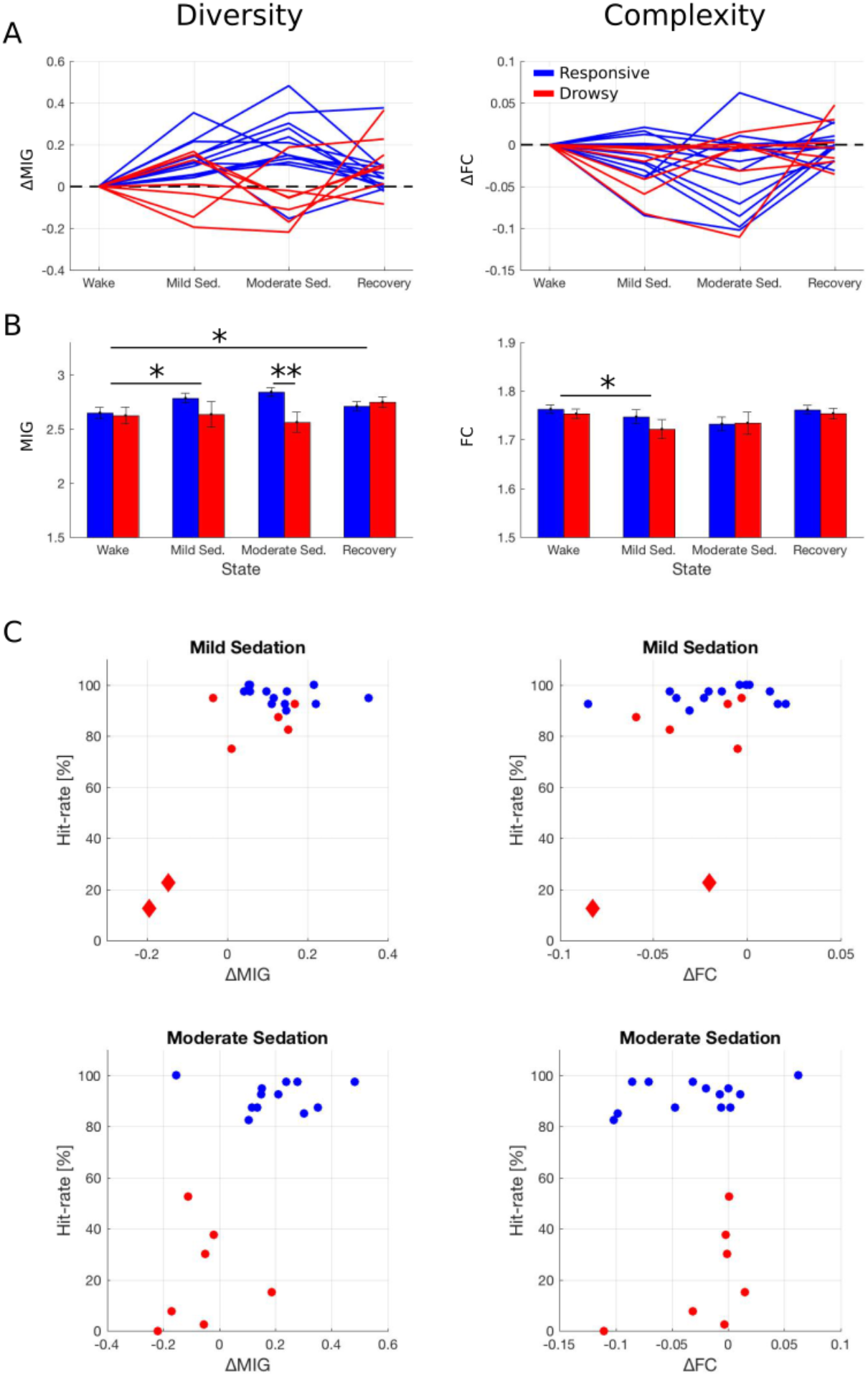
Diversity and complexity of the EEG signal across consciousness states. A) Data of individual subjects from both groups. Data of each were normalized by subtracting the baseline (wake) value to better depict profiles of changes. B) The same data presented as mean ± SEM without normalization. Results of between-states and between-groups comparisons depicted as: * - p < 0.05; ** - p < 0.01. C) Individual hit-rates from the auditory detection task plotted against changes in diversity/complexity measures (with respect to baseline, as in A). Data of two subjects who became unresponsive already during mild sedation state are plotted as red diamonds (compare to Fig. 1).

**Figure 4.**
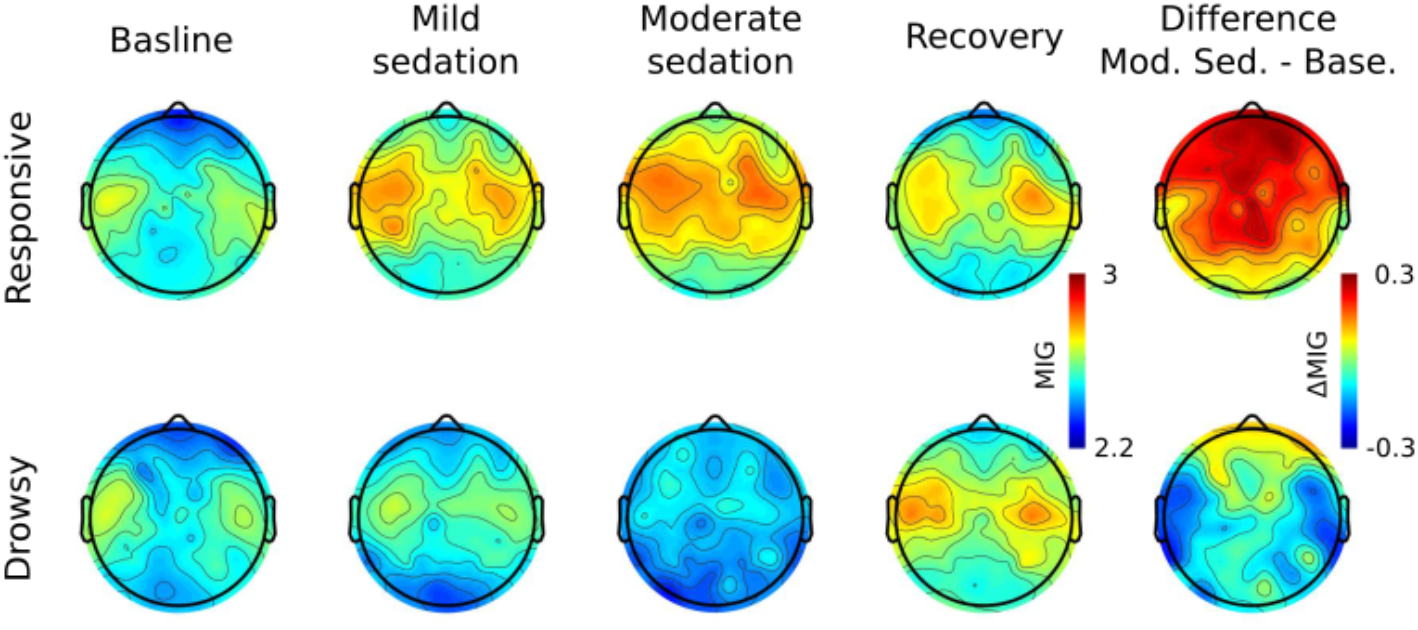
Topographic maps of MIG for *responsive* and *drowsy* groups.

Next we analyzed the Fluctuation Complexity (FC), which is a measures of complexity understood as a state intermediate between an order and disorder. We found a significant effect of *state* (F(3) = 4.46, p = 0.007; BF_10_ = 9.56), but no effect of *group* (F(1) = 0.36, p = 0.55; BF_10_ = 0.50) and no *group x state* interaction (F(3) = 0.74, p = 0.53; BF_10_ = 0.24). Subsequent post-hoc tests indicate that, with respect to baseline wakefulness, FC decreases during mild sedation (t(19) = 3.20, p = 0.028; BF_10_ = 9.65). Comparison of wake with moderate sedation is inconclusive (t(19) = 2.57, p = 0.092; BF_10_ = 3.08), as *p* value reaches a trend level only but BF_10_ indicates evidence for a difference between states. All other comparisons between states were not significant (all p > 0.10; all BF_10_ < 3).

Further, we report results of three additional analyses. First, inspection of the hit-rates profiles suggests that 2 subjects became unresponsive already during the mild sedation state (both had hit-rate < 25%; **Fig. 2**). Plotting hit-rates against changes in MIG demonstrates that exactly these two subjects exhibit a robust decrease in MIG already during mild-sedation (**Fig. 3C**). While statistical analysis cannot be conducted on such a small subsample of 2 subjects, this observation provides an additional, qualitative piece of evidence that MIG is closely related to responsiveness. However, similar scatter-plots created for FC clearly show that FC does not differentiate between responsive and drowsy groups in neither state.

Second, we interpret the difference between groups in MIG during moderate sedation as related to behavioral responsiveness, but the groups differ also in terms of propofol concentration in blood. Thus, to further support our interpretation we compared MIG between groups with propofol concentration included as a covariate. We found a significant difference between groups in MIG (F(1) = 9.53, p = 0.007) but the effect of a propofol concentration was not significant (F(1) = 0.48, p=0.49). Therefore, it is unlikely that the between-groups difference in anesthetic concentration can account for the difference in diversity.

Third, for consistency with several previous studies (e.g. Schartner et al., 2015, 2017, 2017b; Farnes et al., 2019) we applied the Lemepl-Ziv algorithm to estimate signal diversity in our data. In summary, single-channel Lempel-Ziv (LZs) exhibited a pattern of changes which is qualitatively similar to MIG (**Fig. S1–S2**). Results of the Lempel-Ziv analysis can be found in the supplementary materials.

### Relation between diversity/complexity and spectral power

To elucidate a relation between signal diversity/complexity and the spectral content of the analyzed signals, we conducted an analysis of spectral power in four classic EEG frequency bands: delta (1-4 Hz), theta (4-8 Hz), alpha (8-15 Hz), and beta (15-25 Hz). Investigating changes in these bands across states and between the subgroups we found an increase of delta band power during moderate sedation, which was more pronounced in the drowsy than in the responsive group. There was also a robust increase of alpha and beta band power during sedation, but these bands did not distinguish between the subgroups (**Fig. 5**). Detailed statistics can be found in the Supplementary Materials.

**Figure 5.**
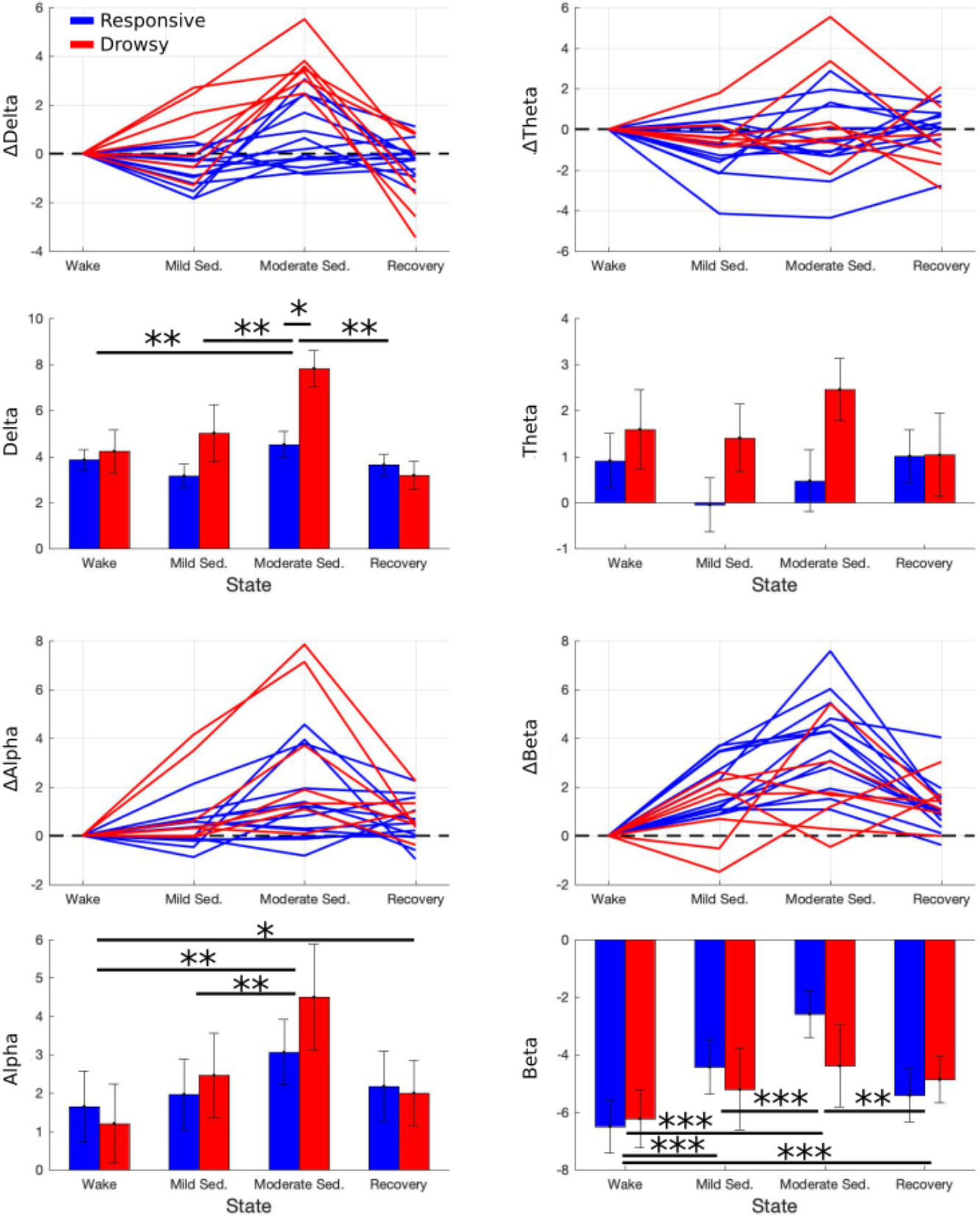
Results of the spectral power analysis in four frequency bands. Data presented in the same way as in Fig. 1. Please note that power estimates were transformed logarithmically and presented as 10log^10^(μV^2^/Hz). Results of between-states and between-groups comparisons depicted as: * - p < 0.05; ** - p < 0.01; *** - p < 0.001.

Next, we correlated measures of diversity and complexity with estimates of the spectral power. Two types of correlations were done, first between diversity/complexity and power at wakeful baseline, and second, between changes in both diversity/complexity and power (i.e. moderate sedation - baseline). Finally, we calculated the Area Under the Receiver Operating Curve (AUC) as an indicator of ability of each measure to distinguish between *responsive* and *drowsy* subgroups. First, we found that high signal diversity is robustly associated with high beta band power (**Table 1**) However, change in MIG dissociated responsive and drowsy groups better than change in the beta band power, which can be seen in the scatter plot (**Fig. 6**) and is confirmed by the AUC analysis (**Table 2**). Second, high signal diversity was also associated with low power of the delta band (also theta and alpha for the Lempel-Ziv diversity). The AUC analysis indicates that out of 8 tested measures change in the delta power is the best predictor when distinguishing both subgroups (AUC = 0.96), and MIG is the second best measure (AUC = 0.87).

**Figure 6.**
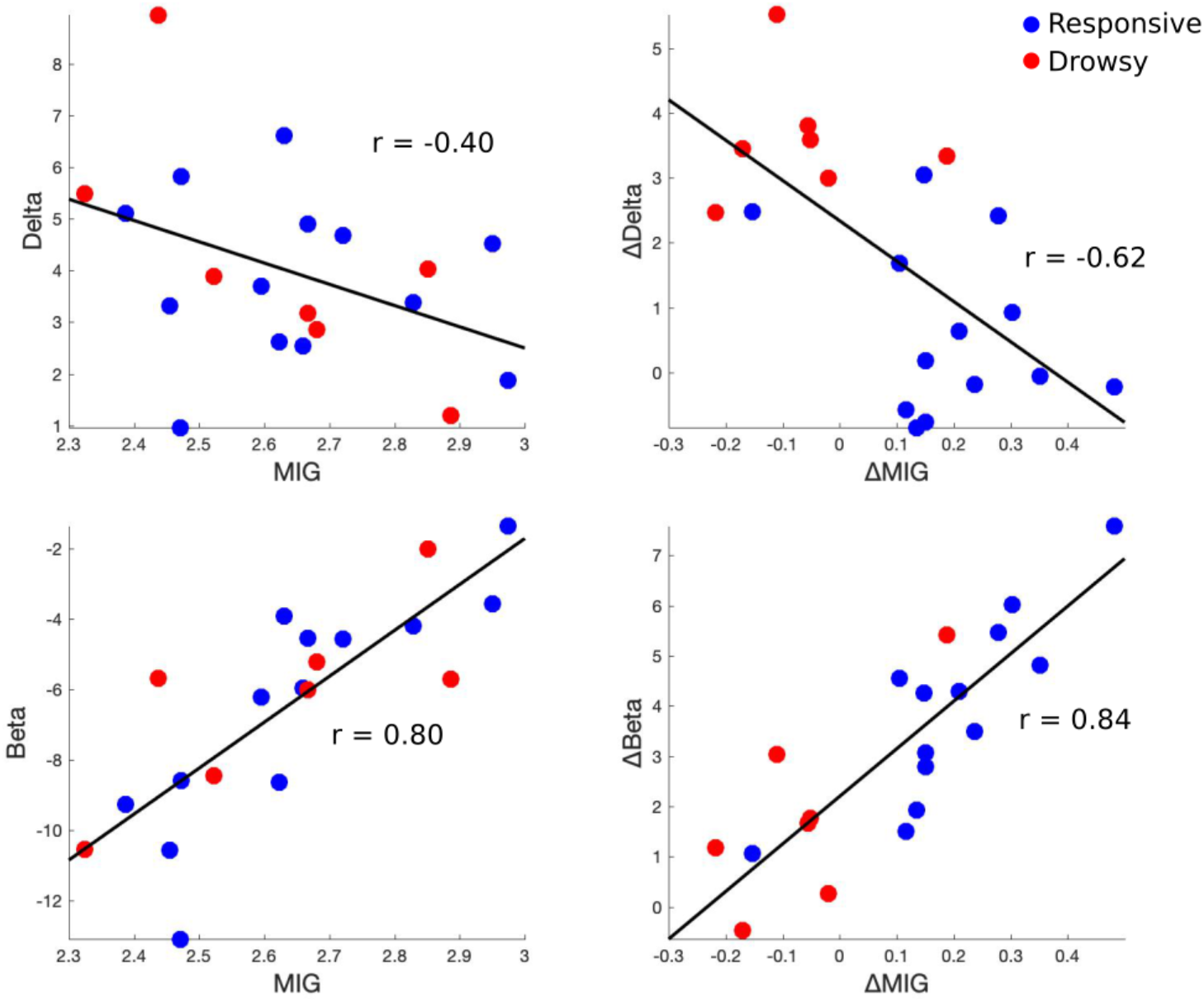
Scatter plots showing a relation between spectral power (delta and beta bands) and signal diversity (MIG). Correlations at baseline wakefulness (left column) and correlations between changes (i.e. baseline wake – moderate sedation; right column) are shown. Black lines indicate best linear fit.

**Table 1.**
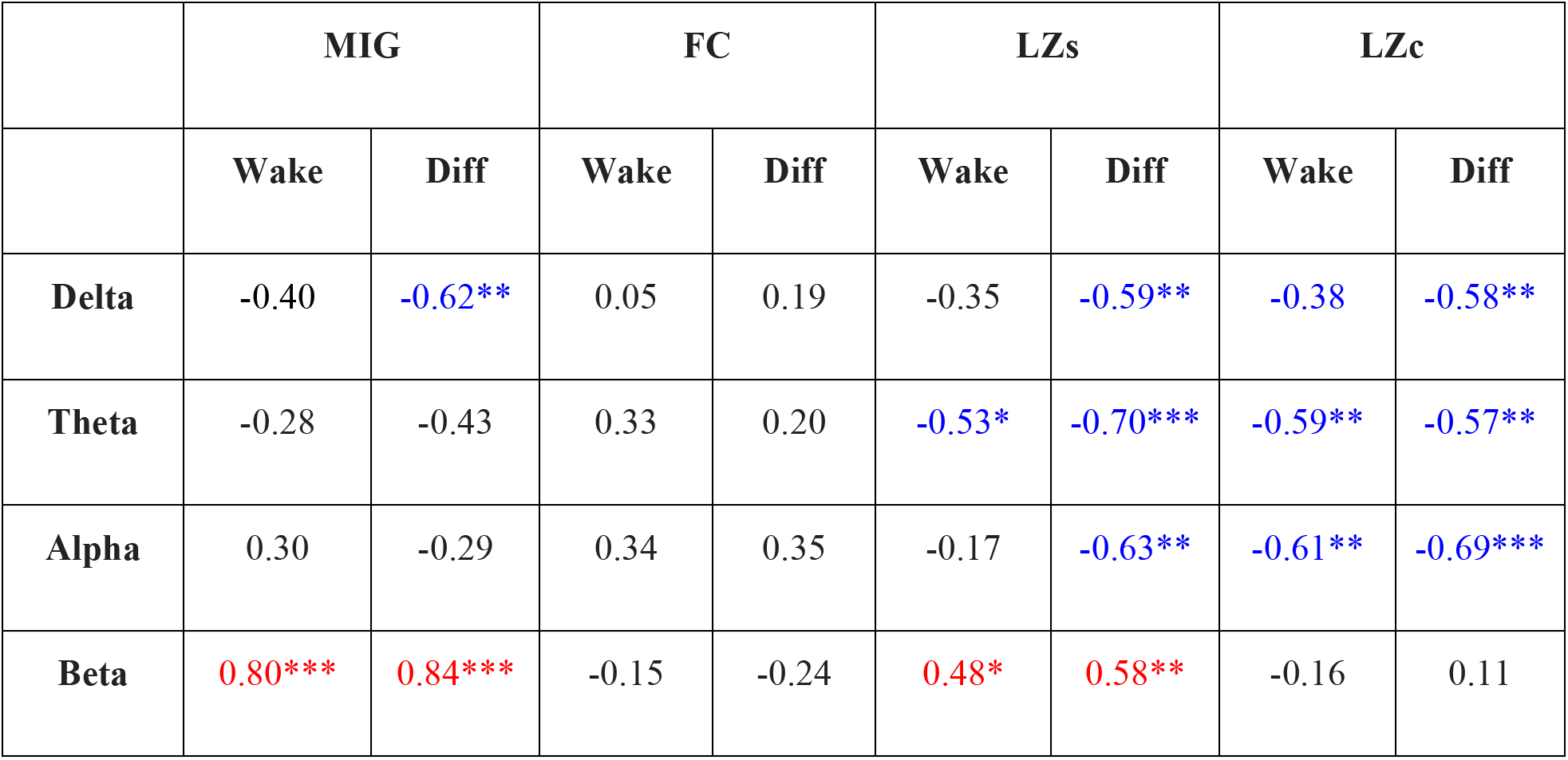
Pearson coefficients (*r*) for correlations between signal power in four bands and diversity/complexity measures. For each measures correlations at wakeful baseline (left column) and correlations between changes (i.e. wake baseline – moderate sedation; right column) are shown. Significance depicted as * - p < 0.05; ** - p < 0.01; *** - p < 0.001. Significant positive correlations are indicated in red, whereas significant negative correlation in blue.

|  | MIG |  | FC |  | LZs |  | LZc |  |
| --- | --- | --- | --- | --- | --- | --- | --- | --- |
|  | Wake | Diff | Wake | Diff | Wake | Diff | Wake | Diff |
| **Delta** | -0.40 | -0.62** | 0.05 | 0.19 | -0.35 | -0.59** | -0.38 | -0.58** |
| **Theta** | -0.28 | -0.43 | 0.33 | 0.20 | -0.53* | -0.70*** | -0.59** | -0.57** |
| **Alpha** | 0.30 | -0.29 | 0.34 | 0.35 | -0.17 | -0.63** | -0.61** | -0.69*** |
| **Beta** | 0.80*** | 0.84*** | -0.15 | -0.24 | 0.48* | 0.58** | -0.16 | 0.11 |

**Table 2.** Results of the receiver operating characteristic curve analysis. Area under the curve (AUC) was calculated for each measure to estimate its performance in classification of the *responsive* and *drowsy* subgroups during moderate sedation.

| | ΔMIG | ΔFC | ΔLZs | ΔLZc | ΔDelta | ΔTheta | ΔAlpha | ΔBeta |
| --- | --- | --- | --- | --- | --- | --- | --- | --- |
| **AUC** | 0.8791 | 0.6044 | 0.7802 | 0.7363 | 0.9670 | 0.6593 | 0.6813 | 0.7912 |

## Discussion

Measures of neuronal diversity have been recently established as robust markers of the global states of consciousness (Casali et al., 2013; review: Storm et al., 2017). Recently a body of evidence has started to emerged suggesting these measures are indeed related to consciousness *per se*, or rather than to some general and unspecific effects occurring during transitions between conscious and unconscious states (Sarasso et al., 2015; Schartner et al., 2017b; Farnes et al., 2019; Schwartzman et al., 2019). Here we aimed to address this question in the context of anesthesia by investigating EEG signal diversity and complexity during propofol sedation. Importantly, despite receiving a similar dose of propofol all tested subjects could be divided into two subgroups: *responsive*, who remained awake throughout the experiment, and *drowsy*, who became unresponsive and unconscious. We hypothesized that the loss of responsiveness during sedation will be related to a decrease of EEG diversity. This was indeed confirmed, as 6/7 subjects from the *drowsy* group exhibited a drop in diversity upon loss of responsiveness. However, the unexpected finding is that EEG diversity increased in 12/13 subjects who were sedated but remained responsive. Such a qualitative difference - an increase in diversity in the responsive group and a decrease in the unresponsive group - makes it a robust index of responsiveness and, presumably, consciousness.

Several recent studies demonstrated that general anesthesia is accompanied by a marked decrease of neuronal diversity as estimated by the Lempel-Ziv algorithm (Sarasso et al., 2015; Schartner et al., 2015). Further evidence for simplification and reduced repertoire of brain states during anesthesia was provided by studies characterizing brain signals in terms of functional interactions (Bola et al., 2018), long-range temporal correlations (Krzemiński et al., 2017; Tagliazucchi et al., 2016), or using dynamic systems theory measures (Alonso et al., 2014; Solovey et al., 2015; Tajima et al., 2015). Our work adds to the literature by demonstrating this effect using “minimal” contrasts between states and groups. By minimal we mean that, first, we detected a robust decrease in diversity upon loss of responsiveness even though the doses of propofol were low (e.g. in comparison to Sarasso et al., 2015 or Schartner et al., 2015). Second, in our analysis we included also sedated but responsive subjects, who therefore constitute an important “control group” missing from previous experiments. Lack of a diversity decrease in this group further strengthens our conclusion that EEG diversity is closely related to the ability to respond to external stimuli.

Whereas the diversity decrease in the drowsy group during moderate sedation is in line with the published results, an increase in the sedated but responsive group is a surprising finding. Further, during mild sedation and recovery - states when all subjects were sedated but responsive - diversity increase was indeed observed in all subjects, irrespective of the subgroup (importantly, propofol was still detected in blood samples during recovery; see: **Fig. 2**). To our knowledge, such an effect has not been reported so far in the context of propofol sedation, even though the state of mild sedation has been included in some of the previous studies. For instance, Schartner et al. (2015) found a decrease in EEG Lempel-Ziv diversity during mild propofol sedation when subjects were partially conscious (Ramsay scale score = 3). However, in the study of Schartner and colleagues the average concentration of propofol in blood during mild sedation was 1.9 ± 0.52 mcg/mL, whereas here it was 0.44 ± 0.04 mcg/mL and 0.90 ± 0.06 mcg/mL during mild and moderate sedation, respectively. Further, Huang et al. (2018) investigated temporal dynamics of BOLD signals with the targeted propofol concentration of 1.3 mcg/mL, whereas Liu et al. (2018) analyzed entropy of BOLD signals with a targeted propofol level of 0.98 mcg/mL. In neither study an effect, which could be interpreted as an increase of diversity or variability of brain activity has been found, which might be attributed to lower sensitivity of fMRI (in comparison to EEG).

However, an increase in EEG signal diversity has been already observed, specifically during psychedelic states induced by LSD or ketamine (Schartner et al., 2017b; Farnes et al., 2019; Wang et al., 2017; Tagliazucchi et al., 2014), or by stroboscopic light stimulation (Schwartzman et al., 2019). Results of these studies suggest two conclusions. First, not only decreases, but also increases of signal diversity with respect to the values observed during resting wakefulness, are in principle possible. Our finding adds to this body of evidence by showing this effect is not specific to the psychedelic states only. Second, brain signal diversity might reflect the diversity of subjective experience. Indeed, in the study of Schartner et al. (2017b) the intensity of psychedelic experience induced by LSD, which subjects self-assessed with a questionnaire, was correlated with an increase in signal diversity. This has been also suggested by experiments showing that signal diversity is higher in response to meaningful visual stimuli, which evoke a range of rich experiences in subjects perceiving them, and lower when subjects perceive stimuli without meaning, which do not result in temporally differentiated experiences (Boly et al., 2015, Mensen et al., 2017, 2018; but see Bola et al., 2018 for a different result in the auditory modality). In the present study we are unable to evaluate experience of sedated but responsive subjects, so we cannot conclude whether an increase in EEG diversity is somehow related to changes in their phenomenology. Therefore, the relation between brain signal diversity and different dimensions of subjective experience (Baynes et al., 2016; Jonkisz et al., 2017) remains one of the most important aspects to investigate.

What is the neurophysiological mechanism behind changes in EEG diversity? Signal diversity and complexity measures can be treated as composite measures reflecting various aspects of the signal. Therefore, to shed some light on this question we analyzed changes in power of the classic EEG frequency bands and related them to changes in EEG diversity/complexity. First, we found reliable increases in delta, alpha and beta power during sedation, but only the magnitude of the delta power increase differentiated the *responsive* and *drowsy* subgroups. An increase of slow-wave (delta band) activity is among most widely studied and most reliable indicators of loss of consciousness, during both sleep and anesthesia (Breshears et al., 2010; Murphy et al., 2011; Purdon et al., 2013). Indeed, in the analyzed data the delta power increase was the most sensitive index of becoming unresponsive to auditory stimuli. Further, changes in the alpha band have also been reported during propofol sedation. Specifically, alpha oscillations shift from the occipito-parietal region to the fronto-central area (a phenomenon called “alpha anteriorization”) and this might result in a more regular and less diverse EEG signal (EEG: Murphy et al., 2011; Supp et al., 2011; Purdon et al., 2013; modeling: Ching et al., 2010). Finally, low anesthetic doses have been previously reported to induce an increase in the beta/gamma bands amplitude (EEG: Gugino et al., 2001; modeling: McCarthy et al., 2008). This effect is considered to represent an initial paradoxical excitation caused by an anesthetic and might potentially result in a diversity increase. Therefore, the pattern of spectral power changes observed in our data is in line with the literature.

Further, to formally assess a relation between spectral power and diversity/complexity we calculated correlations between these measures. We show that greater power of delta band (and to some extent also theta and alpha) is related to lower diversity, whereas greater power of the beta band is related to greater diversity. This observation is in line with another recent study, which also found a strong positive correlation between beta power and Lempel-Ziv diversity (LZs; Schwartzman et al., 2019). Further, Wang et al. (2017) demonstrated a decrease of EEG diversity during propofol sedation, and an increase of EEG diversity during ketamine sedation. While they did not relate diversity measures to spectral power directly, they did report that propofol sedation results in a more pronounced delta power increase, whereas ketamine sedation in a robust gamma power increase. Therefore, one can infer that in the Wang et al. (2017) study diversity would also be negatively related to low-frequency power, and positively to high-frequency power.

In the present analysis diversity of EEG signals was estimated by the Mean Information Gain (MIG), which is an information theory measure (Bates and Shepard, 1993). However, multiple previous studies established the Lemepl-Ziv algorithm as a robust correlate of the global conscious states (e.g. Casali et al., 2013; Schartner et al., 2015). Therefore, we also calculated the Lemepl-Ziv diversity to compare it with MIG (results reported in the supplementary materials). While both measures behaved in a qualitatively similar way, MIG seems to distinguish responsive and drowsy groups more robustly, and only MIG increased significantly during mild sedation and recovery. Greater sensitivity of MIG might be explained by the fact that, for consistency with previous studies, MIG was applied to the EEG signal (Wang et al., 2017), whereas the Lempel-Ziv algorithm was applied to the signal envelopes (Schartner et al., 2015, 2017; Bola et al., 2018). Further, Lempel-Ziv was used in two versions, to estimate diversity of either temporal (LZs) or spatio-temporal (LZc) patterns. Our results demonstrate that only LZs, but not LZc, was sensitive enough to distinguish between responsive and drowsy groups. This suggests that temporal (rather than spatial) diversity of brain activity is the key aspect and should be the focus of future studies.

Nevertheless, it is important to keep in mind that both MIG and Lempel-Ziv are measures of signal *diversity*, defined as randomness or entropy of the data, and not *complexity*, understood as a state intermediate between order and randomness. Therefore, in our study a measure of complexity, namely Fluctuation Complexity (FC), was also included. FC has been proved capable of robustly differentiating states of anesthesia and wakefulness (Wang et al., 2017). However, FC decreased only during mild and not during moderate sedation, and it did not distinguish responsive and drowsy subjects (**Fig. 3**). Lack of a reliable complexity decrease during loss of responsiveness might be related to the fact, that moving from low diversity to higher diversity causes an increase in complexity (i.e. moving towards the peak of the inverted “U” shape, see: **Fig. 1**). Therefore, our analysis suggests that subjects’ behavioral state during propofol sedation is better represented by diversity than complexity of the EEG signal.

In summary, by comparing responsive and unresponsive subjects during propofol sedation we provide further evidence that EEG signal diversity is a promising correlate of responsiveness. Key questions which remain to be addressed concern the physiological mechanism behind signal diversity changes, and the relation between signal diversity and various aspects of phenomenology.

## Supplementary Materials

### Supplementary Results

#### Lemepl-Ziv diversity

In the present study we focused on two information theory measures - MIG, which is a measure of diversity, and FC, which is a measure of complexity. However, for consistency with multiple previous studies which used the Lempel-Ziv algorithm we also calculated two version of Lempel-Ziv diversity on our data. The implementation used here is the same as in Schartner et al. (2017a).

First, we investigated temporal diversity, denoted here as LZs (**Fig. S1–2**). Statistical analysis of LZs indicates no main effect of *group* (F(1) = 1.9, p = 0.18; BF_10_ = 0.75) and no effect of *state* (F(3) = 1.48, p = 0.22; BF_10_ = 1.05), but a significant *group x state* interaction was found (F(3) = 3.27, p = 0.028; BF_10_ = 3.0). Subsequent post-hoc tests indicate no differences between states (all p > 0.05). However, comparisons between groups indicate that the drowsy group exhibits lower LZs during moderate sedation (t(18) = 3.32, p = 0.004; BF_10_ = 10.4) but not during other states (all p > 0.10; all 0.3 < BF_10_ < 1).

Second, we investigated spatio-temporal diversity, denoted here as LZc. There was no main effect of *group* (F(1) = 0.35, p = 0.56; BF_10_ = 0.56), a significant effect of *state* (F(3) = 4.76, p = 0.005; BF_10_ = 7.24), and no *group x state* interaction (F(3) = 0.74, p = 0.53; BF_10_ = 1.0). However, subsequent post-hoc tests indicate a significant difference only between moderate sedation and recovery (t(19) = 3.15, p = 0.031; BF_10_ = 8.82), with all other comparisons not reaching significance threshold (all p > 0.10; all 0.3 < BF_10_ < 1).

#### Spectral power

Statistical analysis of the delta-band power indicates no main effect of *group* (F(1) = 2.2, p = 0.15; BF_10_ = 0.90) but a significant effect of a *state* (F(3) = 27.04, p < 0.001; BF_10_ > 1000) and a significant *group x state* interaction was found (F(3) = 12.84, p < 0.001; BF_10_ > 100). Post-hoc tests reveal an increase in delta power during moderate sedation with respect to all other states (baseline: t(19) = 4.07, p = 0.004; BF_10_ = 53; mild sedation: t(19) = 4.67, p < 0.001; BF_10_ = 177; recovery: t(19) = 4.18, p = 0.003; BF_10_ = 66), but other comparisons were not significant (p > 0.1). Further, the delta power increase during moderate sedation was more pronounced in the drowsy group than in the responsive group (t(1) = 3.46, p = 0.003; BF_10_ = 13.23).

Theta power did not exhibit a *group* (F(1) = 1.38, p = 0.25; BF_10_ = 0.81) or *state* effect (F(3) = 1.34, p = 0.26; BF_10_ = 0.27), nor a significant *group x state* interaction (F(3) = 2.20, p = 0.098; BF_10_ = 0.92). Therefore, post-hoc comparisons were not conducted.

Alpha power did not exhibit a *group* effect (F(1) = 0.05, p = 0.81; BF_10_ = 0.60), but there was a significant effect of the *state* (F(3) = 13.31, p < 0.001; BF_10_ > 1000). No significant *group x state* interaction was found (F(3) = 2.37, p = 0.081; BF_10_ = 0.78). Subsequent post-hoc comparisons uncovered that moderate sedation differed from wakeful baseline (t(1) = 3.88, p = 0.006; BF_10_ = 36) and mild sedation (t(1) = 3.79, p = 0.007; BF_10_ = 30), and additionally there was a difference between wakeful baseline and recovery (t(1) = 3.14, p = 0.032; BF_10_ = 8.7).

Beta power did not exhibit a *group* effect (F(1) = 0.09, p = 0.76; BF_10_ = 0.55), but there was a significant effect of the *state* (F(3) = 18.19, p < 0.001; BF_10_ > 1000) and a significant *group x state* interaction (F(3) = 3.75, p = 0.016; BF_10_ = 4.51). Post-hoc tests indicate significant differences between most of the pairs of states, specifically beta power increased during moderate sedation with respect to wakeful baseline (t(1) = 6.79, p < 0.001; BF_10_ > 1000), mild sedation (t(1) = 3.31, p < 0.022; BF_10_ = 12), and recovery (t(1) = 4.22, p = 0.003; BF_10_ = 71). Further beta power was higher during mild sedation than wakeful baseline (t(1) = 5.55, p < 0.001; BF_10_ > 1000), and during recovery in comparison to baseline (t(1) = 5.25, p < 0.001; BF_10_ = 567). However, comparisons between the sub-groups did not show differences in any pair of the states (all p > 0.1).

### Supplementary Figures

**Figure S1.**
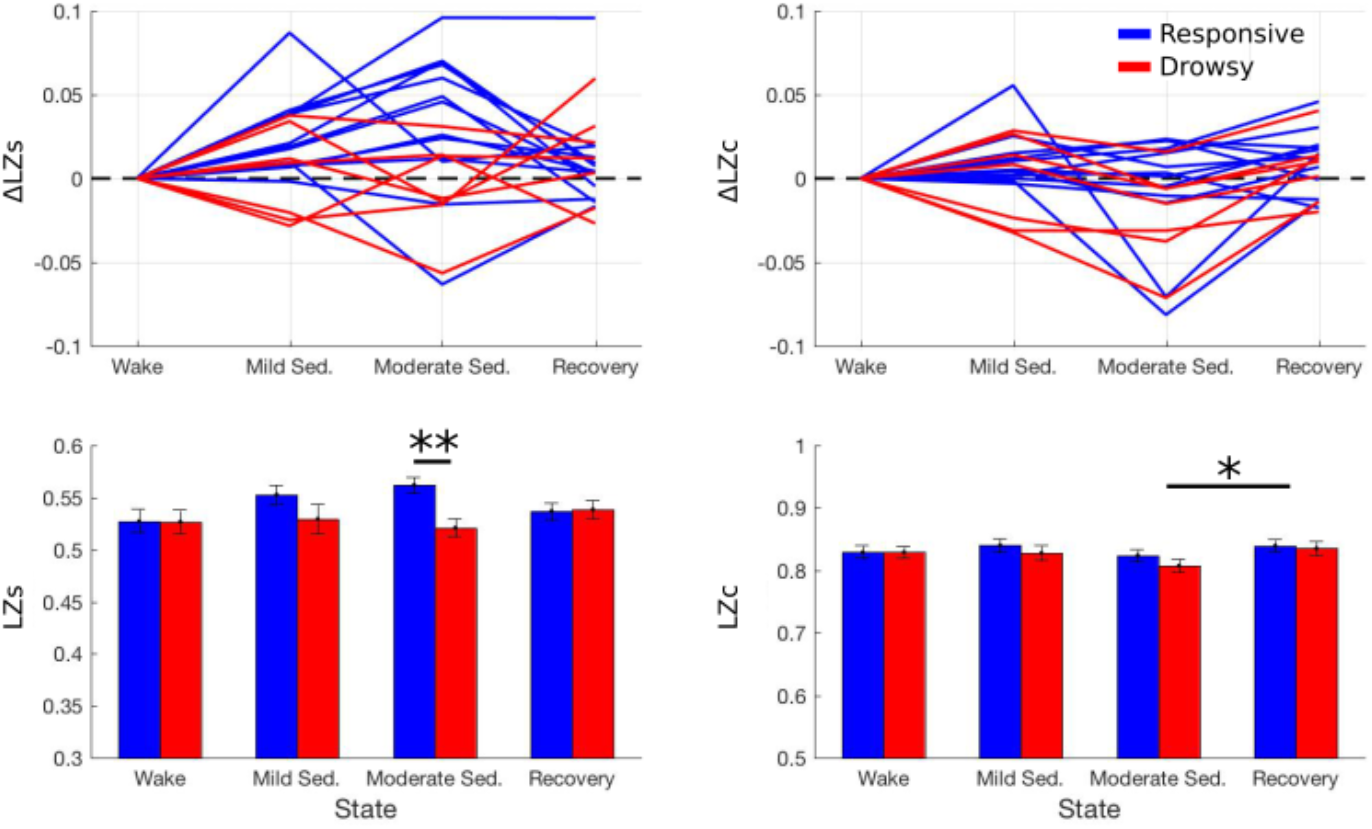
Two versions of Lemepl-Ziv, measuring either temporal (LZs) or spatio-temporal (LZc) diversity of EEG data. A) Data of individual subjects from both groups. To depict profiles of changes data of each subject were normalized by subtracting the baseline (wake) value. B) The same data presented as mean ± SEM without normalization. Results of between-states and between-groups comparisons depicted as: * - p < 0.05; ** - p < 0.01.

**Figure S2.**
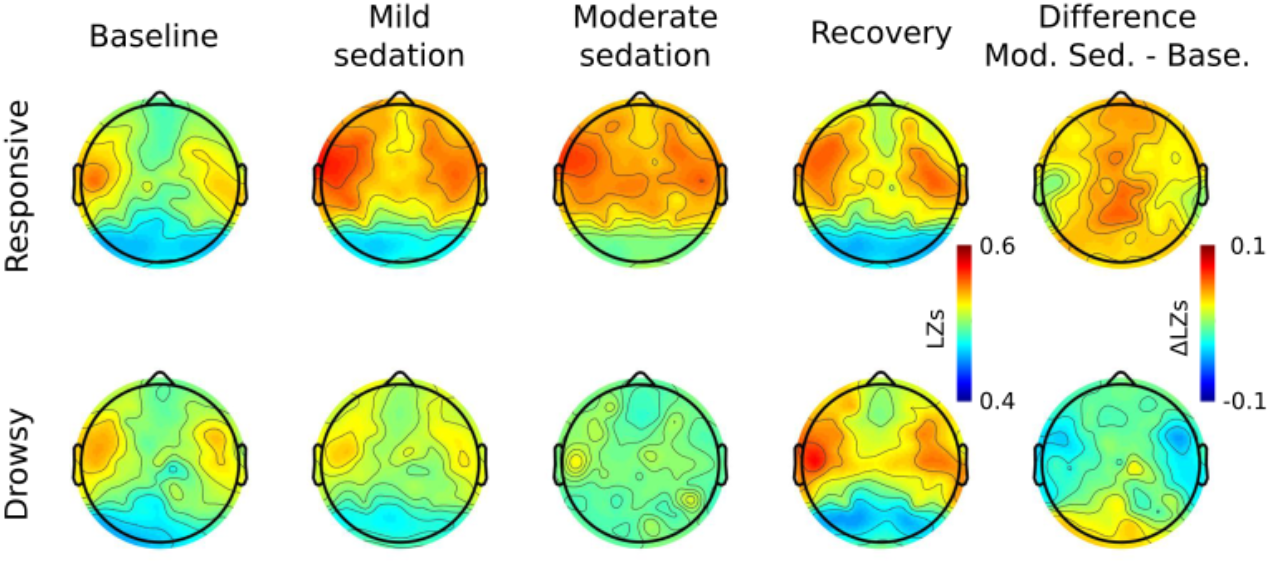
Topographic maps of LZs for *responsive* and *drowsy* groups (compare to Fig. 4).

## Acknowledgments

This study was funded by Sonata grant from the National Science Centre Poland (2015/17/D/HS6/00269) and Iuventus Plus grant from the Polish Ministry of Science and Higher Education (082/IP3/2016/74). MB was additionally supported by a stipend from the Polish Ministry of Science and Higher Education (555/STYP/11/2016). We thank Srivas Chennu for his helpful suggestions regarding the analysis.

Author Contributions
Conceived the study: MB; Analyzed data: PO, MB; Contributed analysis methods: MP; Interpreted data: MB, AM; Wrote the paper: MB. Revised the paper: MB, AM.

